# Stem rot affects the structure of rhizosphere microbiome in Berseem Clover (*Trifolium alexandrinum*)

**DOI:** 10.1101/2024.03.13.584667

**Authors:** Salma Mukhtar, Zain Ahmad, Noor Khan, Michael John, Dalaq Aiysha

**Affiliations:** Department of Plant Biology, Rutgers University, New Jersey, USA; Biochemistry and Molecular Biology, University of Agriculture, Faisalabad, Pakistan; University of California, Los Angeles, USA; Stockbridge School of Agriculture, University of Massachusetts, USA; Department of Microbiology and Molecular Genetics, University of the Punjab, Pakistan

**Keywords:** Stem rot, Rhizosphere microbiome, Berseem Clover, *Sclerotinia sclerotiorum*

## Abstract

Rhizosphere microbiome plays an essential role in maintaining plant health and productivity. Fungal and bacterial diseases may affect the rhizosphere-associated microbial communities and overall structure of plant microbiome. Here, we studied the effect of stem rot of berseem clover on the bacterial and fungal communities associated with the rhizosphere. We analyzed the rhizosphere-associated bacterial and fungal microbiome from healthy and infected berseem clover collected from three sampling sites by using 16S rRNA and ITS based Illumina sequencing metabarcoding approach. Microbiome analysis showed that healthy plants had higher bacterial and fungal diversity as compared to stem rot infected plants. At the genus level, bacterial genera *Rhizobium* and *Comamonas* were more abundant in healthy plants while *Pantoea* was more abundant in infected plants and fungal genera *Sclerotinia*, *Fusarium* and *Cladorrhinum* were more abundant in infected plants while *Microdochium* and *Cladosporium* were distinctively abundant in healthy Berseem. Functional characterization of bacterial and fungal microbiomes revealed that bacterial communities from infected plants showed more abundance of bacteria with functions replication and repair, enzyme families and biosynthesis of other secondary metabolites as compared to healthy plant microbiome and decreased in fungal groups including arbuscular mycorrhiza and soil saprotrophs and an increase in plant saprotrophs and fungal parasite-plant pathogens. This study provides comprehensive information about the structure and composition of bacterial and fungal communities associated with the berseem clover rhizosphere that could be utilized for future research on the control of stem rot of berseem clover.

## Introduction

Berseem clover (*Trifolium alexandrinum* L.) is an economically important crop in the world, with nearly 90% of smallholding farmers cultivating it in different climates and soil types. It is a winter crop cultivated in different regions of Pakistan. It is a multipurpose crop that is used for cover cropping, non-bloating forage production, green manure, and honey production (Saeed et al. 2011). It can be cultivated with the rotation of other crops. Its production is low (3354 t/ha) in Pakistan as compared to other countries. Poor agronomic methods, unavailability of well-adapted seed varieties and timely input are some important factors that contribute to low production (Anwar et al. 2012; Tufail et al. 2020).

Berseem clover is affected by various organisms including fungi, viruses and nematodes. Some common fungal diseases in berseem are stem rot (*Sclerotinia sclerotiorum*), root rot (*Rhizoctonia solani, Fusarium moniliforme, and Sclerotinia bataticola*), clover scorch (*Kabatiella caulivora*) and damping off (*Pythium spinosum*) (Singh et al., 2020).

Stem rot of berseem is caused by a fungus known as *Sclerotinia sclerotiorum*. This fungal disease is widely distributed and infects about 400 species of berseem globally (Saira et al. 2017). The disease is reported to cause more than $270 million in losses. In Pakistan, the disease is infecting all cultivars of Berseem and causing heavy losses. The Sargodha, Kasur, Faisalabad and Okara districts are the main Berseem cultivation areas of Punjab, Pakistan. This disease is subject to significant crop losses and drastically reduced growth. An integrated disease management approach employing biological and chemical control was adopted to manage this wide-spreading fungal pathogen (Atri et al. 2021).

Microbial communities associated with the rhizosphere play an important role in plant health and soil fertility under abiotic and biotic stress conditions. Root cells produce different exudates that can be utilized by rhizosphere microorganisms (Zhao et al., 2016; Santhanam et al., 2017; Banerjee et al., 2018). These microbes enhance plant growth by increasing the availability of nutrients, nitrogen fixation, production of phytohormones and siderophore and control many fungal and bacterial diseases (Chen et al. 2020; Mukhtar et al. 2020; Qu et al. 2020). Rhizobia and other bacteria can be used as biocontrol agents and biofertilizers to improve the clover yield. Several bacterial genera including *Bacillus, Enterobacter, Agrobacterium, Acinetobacter, Paenibacillus, Micromonospora, Pantoea,* and *Pseudomonas* have also been isolated from rhizosphere of different legume plants (Zgadzaj et al. 2015; Leite et al. 2017; Liu et al. 2017).

Plant genotype, abiotic and biotic stresses influence the structure and composition of the rhizosphere microbiome (Mukhtar et al. 2021; Shi et al. 2021; Oyserman et al. 2022). Stem or root fungal diseases are difficult to control because soil borne fungi are usually ubiquitous in nature, so these pathogens often occur as a complex and they can easily survive on infected plant debris or form durable in the soil with or even in the absence of growing hosts and can outgrow or evade plant defenses (Wintera et al. 2014). Therefore, it is very critical to study the relationship between plants and rhizosphere microbiome to understand different fungal and bacterial diseases. Rhizosphere microbial diversity directly affects plant health, stress tolerance, productivity and fruit yield (Kristin and Miranda 2013; Miao et al. 2016; Zhang et al. 2021)

Most previous studies have focused on how the microbial communities are influenced by host genotypes and abiotic factors such as salinity and drought (Rahal and Chekireb 2021; Youseif et al. 2021; Kozieł et al., 2022). We hypothesized that because of a fungal infection, bacterial and fungal communities associated with the rhizosphere of berseem clover were changed. These microorganisms help to control or mediate disease resistance. Here, we evaluated the response of rhizosphere microorganisms by comparative analysis of healthy and infected plants. To the best of our knowledge, this study is the first report on the response of the rhizosphere microbiome of berseem clover to stem rot infection.

## Material and methods

### Soil and plant sampling

Rhizosphere soil samples from healthy and infected berseem clover were collected from 3 different locations in district Sargodha. These locations were chosen with the following geographical details; site 1 was located at 32°41’27“N, 72°13’10”E site 2 was located at 32°45’59“N, 72°33’18”E and site 3 was located at 32°53’37“N, 72°19’41”E. Soil sampling for microbiome analysis was carried out at flowering stage. From each site, six healthy and six infected plants were randomly selected (**Fig. S1**). Rhizospheric soil samples were collected by gently removing the plants and obtaining the soil attached to the roots. At each site, approximately 1 kg soil samples were collected in black sterile polythene bags. These samples were stored at -80 L for further analysis.

### Soil physicochemical parameters

Each soil sample (500 g) was thoroughly mixed and sieved through a pore size of 2 mm. Physical properties (salinity, pH, moisture content and temperature) of soil samples were determined. Soil salinity or electrical conductivity (dS/m) was measured by 1:1 (w/v) soil to water mixture at 25 L (Adviento-Borbe et al. 2006), pH was measured by 1:2 (w/v) soil to water mixture, moisture (%) and texture class were measured by Anderson method (Anderson et al. 1993) and organic matter (C_org_) was calculated by the Walkley-Black method (1934).

### DNA extraction from rhizosphere samples

DNA was extracted from 250 mg rhizosphere soil by using the DNeasy PowerSoil Kit (Qiagen, Hilden, Germany) according to the manufacturer’s instructions. DNA yield was quantified by using the qubit method (Thermo Invitrogen Qubit 4).

### PCR amplification of 16S rRNA and ITS genes and sequencing

The V3-V4 region of the bacterial 16S rRNA gene was amplified by using the primer pair 338F(5’-ACTCCTACGGGAGGCAGCAG-3’) and 806R (5’- GGACTACHVGGGTWTCTAAT-3’). The 5’ end of both primers were tagged with specific barcodes and universal sequencing primers (Sinclair et al. 2015). PCR amplification of 16S rRNA gene was performed as previously described by Caporaso et al. (2011). PCR products with the correct amplicon size were confirmed on 1% gel electrophoresis. Sterile water was used as a template instead of DNA in negative control reactions. In gel electrophoresis of negative reactions, no bands were observed. PCR products were purified by using the QIAquick PCR purification kit (Qiagen, Hilden, Germany).

The ITS1 region of the eukaryotic (fungi) small-subunit rRNA gene was amplified by using primers ITS1f and ITS1r (Taylor et al. 2016). PCR products were confirmed by using 1% gel electrophoresis. Sterile water was used as a negative control for this PCR. Then these PCRproducts were digested by using two restriction enzymes including exonuclease I and antarctic phosphatase at 37 °C for 1 hr. The digested PCR products were again amplified by using sequencing primers with specific barcodes at the 5’ ends. PCR products were confirmed with 1% agarose gel electrophoresis. PCR products were purified by using the QIAquick PCR purification kit (Qiagen, Hilden, Germany).

Amplicons for 16S rRNA and ITS genes were pooled separately in two tubes. The size and quantity of the amplicon for both libraries were assessed using an Agilent 2100 Bioanalyzer (Agilent, USA). Normalized amplicons were pooled together and submitted for sequencing using Miseq v2.2.0 platform (Illumina Inc, San Diego CA).

### Sequence processing and microbiome analysis

Raw amplicon sequences were joined and demultiplexed using qiime2 pipeline (Bolyen et al. 2019). Chimeric and PhiX were filtered out using qiime2-DADA2 (Callahan et al. 2016). Taxonomic classification of reads of 16S rRNA sequences was performed using qiime2 feature classifier (Green Genes 13.8.99) and the primer set 338F/806R. Reads were assigned to operational taxonomic unit (OTUs) with 100% sequence identity (Pedregosa et al. 2011). Classification of ITS OTUs was carried out by using the UNITE database (version 04.02.2020) (UNITE Community 2019). OTUs Alpha-diversity, Shannon and Observed OTUs was analyzed by using the R package “phyloseq” (McMurdie and Holmes 2013) and Faith’s phylogenetic diversity was analyzed by using the R package “picante” (Kembel et al. 2010). Count data were rarefied to 41626 reads. Kruskal-Wallis and Kruskal-Conover post-hoc tests were used to test for significant differences. Reads were normalized using cumulative sum scaling and Bray-Curtis distances and projecting constrained principal coordinate analysis (PCoA) and community homogeneity measured by distances to centroid by using the R package “vegan” (Oksanen et al 2022). Constrained Principal coordinates analyses were computed using BC distances using the R function capscale in the package “vegan” (v2.6-2, Oksanen et al 2022). Differences between group medians were tested using Kruskal-Wallis significance test. All plotting graphs were displayed using the R package ggplot2 (v3.3.6, (Wickham et al. 2022). Permutational multivariate analysis of variances in BC distances was computed using the R function adonis2 in the package “vegan”. The raw reads were deposited into the NCBI Sequence Read Archive (SRA) database under the accession numbers SRR27988892-SRR27988927.

### Function Prediction of bacterial and fungal communities

For bacterial function prediction in healthy and infected plants, the analysis was carried out by using the PICRUSt software (Langille et al. 2013). Based on reporter scores from Z-scores of individual Kos (Kyoto Encyclopedia of Genes and Genomes (KEGG Orthology), differentially enriched KO pathways or modules were identified. Wilcoxon rank sum test was applied to all KOs and corrected for multiple testing by the Benjamin-Hochberg method. The Z-score of each KO was computed separately. The detection threshold of significantly differentiating pathways was set as an absolute value for a reporter score ≥ 1.87. To analyze the functional groups of fungi, FUNGuild database was used to classify OTUs into an ecological guild (Nguyen et al., 2016).

## Results

### Soil physiochemical analysis

Electrical conductivity (EC1:1) of soil samples was almost similar ranging from 1.01 dS/m as observed in HBSC3 soil to 1.11 dS/m in DBSC2 soil, pH of collected soil samples ranged from 7.16 for DBSC2 soil and 7.45 for HBSC1 soil, maximum moisture content (%) 26 was observed in two soil samples i.e., DBSC2 and DBSC3 while minimum 20 observed in HBSC2 soil (**Table 1**). All soils had sandy loam texture with variable organic matter content (g/Kg) ranging from 26.11 in HBSC3 to 33.55 as observed in DBSC1 soil. The maximum phosphorous content (mg/Kg) was 4.11 in DBSC3 and a minimum of 3.16 in HBSC3 soil, and potassium content (mg/Kg) ranged from 0.45 in HBSC1 soil to 0.52 in DBSC2 soil, calcium content (mg/Kg) ranged from 1.30 in HBSC3 soil to 1.35 in DBSC2 soil, magnesium content (mg/Kg) ranged from 1.11 in HBSC2 soil to 1.19 in DBSC2 soil. The highest average nitrate ion content (mg/Kg) was 12.55 in the HBSC2 soil and the lowest was 10.23 in the HBSC3 soil sample (**Table 1**).

**Table 1.** Physico-chemical properties of rhizosphere soil of healthy and infected berseem clover.

| Parameters | HBSC1 | HBSC2 | HBSC3 | DBSC1 | DBSC2 | DBSC3 |
| --- | --- | --- | --- | --- | --- | --- |
| EC<sub>1:1</sub> (dS/m) | 1.09<sup>a</sup> | 1.10<sup>a</sup> | 1.11<sup>a</sup> | 1.04<sup>a</sup> | 1.01<sup>a</sup> | 1.03<sup>a</sup> |
| pH | 7.45<sup>b</sup> | 7.36<sup>b</sup> | 7.38<sup>b</sup> | 7.18<sup>a</sup> | 7.16<sup>a</sup> | 7.21<sup>a</sup> |
| Moisture (%) | 22<sup>a</sup> | 20<sup>a</sup> | 21<sup>a</sup> | 24<sup>ab</sup> | 26<sup>b</sup> | 26<sup>b</sup> |
| Texture class | Sandy loam | Sandy loam | Sandy loam | Sandy loam | Sandy loam | Sandy loam |
| OM (g.Kg<sup>-1</sup>) | 27.7<sup>a</sup> | 24.34<sup>a</sup> | 26.11<sup>a</sup> | 33.55<sup>b</sup> | 30.34<sup>b</sup> | 31.11<sup>b</sup> |
| P (mg.kg<sup>-1</sup>) | 3.21<sup>a</sup> | 3.25<sup>a</sup> | 3.16<sup>a</sup> | 4.02<sup>b</sup> | 3.95<sup>b</sup> | 4.11<sup>b</sup> |
| K (mg.kg<sup>-1</sup>) | 0.45<sup>a</sup> | 0.49<sup>a</sup> | 0.51<sup>a</sup> | 0.49<sup>b</sup> | 0.52<sup>b</sup> | 0.51<sup>b</sup> |
| Ca (mg.kg<sup>-1</sup>) | 1.32<sup>a</sup> | 1.33<sup>a</sup> | 1.30<sup>a</sup> | 1.31<sup>a</sup> | 1.35<sup>a</sup> | 1.34<sup>a</sup> |
| Mg (mg.kg<sup>-1</sup>) | 1.13<sup>a</sup> | 1.11<sup>a</sup> | 1.12<sup>ab</sup> | 1.18<sup>b</sup> | 1.19<sup>b</sup> | 1.16<sup>ab</sup> |
| NO<sup>-3</sup> (mg.kg<sup>-1</sup>) | 10.43<sup>a</sup> | 12.55<sup>b</sup> | 10.23<sup>a</sup> | 12.17<sup>b</sup> | 12.24<sup>b</sup> | 10.29<sup>a</sup> |

Note: EC (Electrical conductivity); OM (Organic matter); P (Phosphorous); K (Potassium); Ca (Calcium); Mg (Magnesium); NO<sup>-3</sup> (Nitrate ion); Alphabets represent statistically different values at 5% level.

### Stem rot affects the structure of berseem rhizosphere microbiome

To see the impact of stem rot on berseem clover rhizosphere microbiome we analyzed the rhizospheric microbial communities associated with infected and healthy plants grown under natural conditions. The results of observed OTUs for bacterial communities identified were higher in healthy plants’ rhizosphere as compared to the infected ones with high dimensionality in site 3 (DBCS3) in comparison to the other two sample collection sites (**Fig. 1A-1C**) but it was completely the opposite in the case of observed OTUs for fungal communities where it was highest in healthy plants’ soil as compared to the infected plants’ soil with high dimensionality in site 3 (HBCS3) soil sample (**Fig. 1D-1F**).

**Fig. 1.**
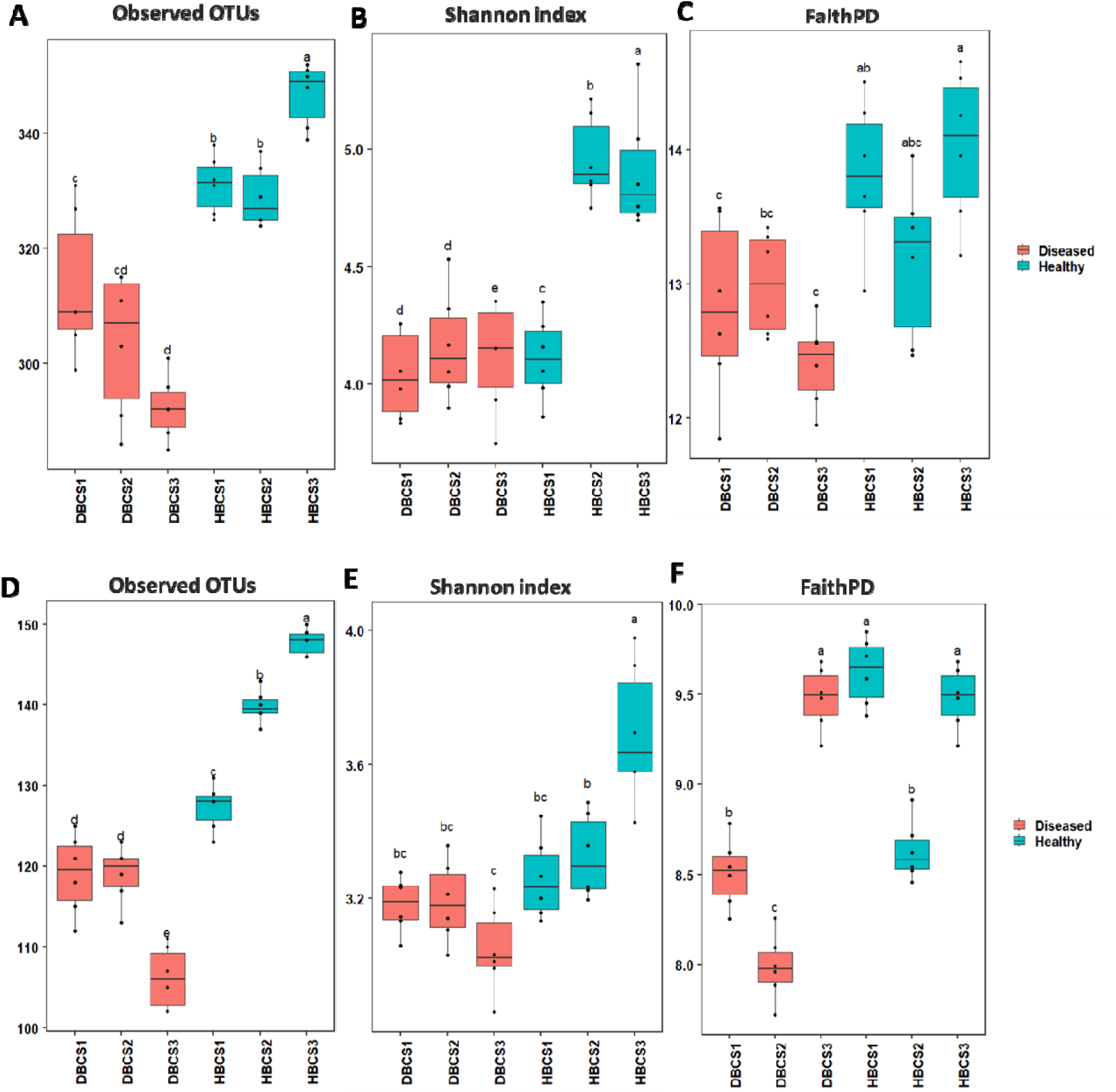
Stem rot influenced the structure of rhizosphere microbiome in Berseem Clover. **A, B and C** showed the boxplots for bacterial Observed OTUs, Shannon’s index, and Faith phylogenetic diversity index, respectively. **C, D and E** showed the fungal bacterial Observed OTUs, Shannon’s index, and Faith phylogenetic diversity index, respectively. The alpha diversity of the healthy berseem microbiome is significantly different from the infected microbiome.

The results of the alpha diversity analysis (Shannon index) showed that the bacterial communities’ evenness and the richness observed in healthy soil samples were significantly different from the infected plant’s rhizosphere. All three sites of the infected plant soil showed similar low diversity in comparison to the two healthy plant soil sites (HBCS2, HBCS3) where site 1 (HBCS1) had lower diversity than the other two sites (**Fig. 1B**). The fungal communities’ evenness and the richness of infected plant rhizosphere were significantly lower than the healthy ones and the highest diversity was observed in the site 3 samples of healthy plants’ rhizosphere (HBCS3) in comparison to all other selected sites (**Fig. 1E**). The microbial community richness in the infected and healthy plants’ rhizosphere was analyzed using the Faith phylogenic diversity index. The bacterial communities showed more richness and diversity in healthy plant rhizosphere highest in the site 3 sample (HBCS3) (**Fig. 1C**) while higher fungal community richness was observed in healthy plant soils highest in the site 1 soil sample (HBCS1) (**Fig. 1F**). The beta diversity as described using constrained principal coordinate analysis of the Berseem rhizosphere microbiome was analyzed using Bray-Curtis distance analysis, which showed that bacterial communities of healthy Berseem rhizosphere were clustered separately as compared to the stem rot affected Berseem rhizosphere. In the healthy Berseem rhizosphere, the bacterial communities found in sample sites 1 and 2 were more similar as compared to the site 3 sample but a similar pattern was not observed in the infected Berseem rhizosphere (**Fig. 2A**). Our results showed that fungal communities of healthy and infected Berseem rhizosphere were dissimilar but clustered together randomly with stability among each sampling site (**Fig. 2B**). Our findings described that stem rot disease has a major impact on the composition and structure of bacterial and fungal communities.

**Fig. 2.**
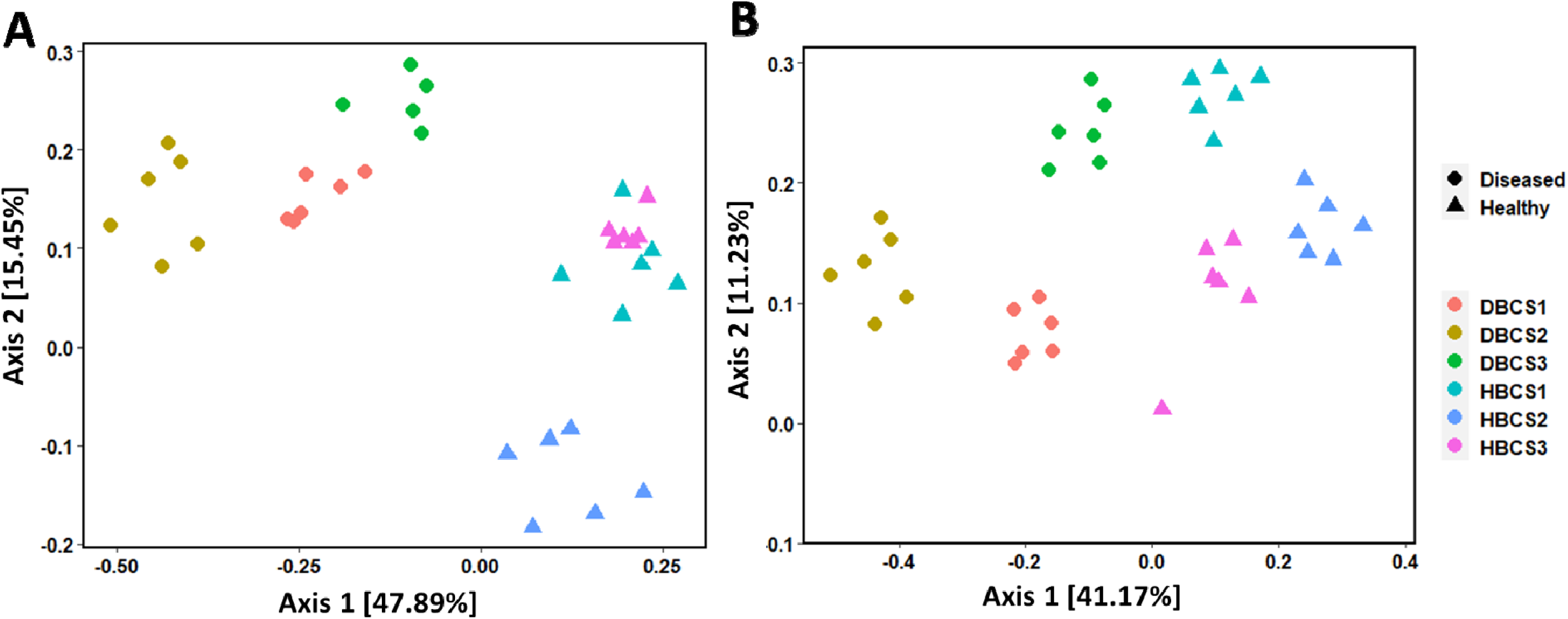
Beta diversity analysis from healthy and infected berseem. **(A)** Constrained principal coordinate analysis (PCoA) showing the impact of stem rot on the rhizosphere bacterial community of berseem. **(B)** Constrained principal coordinate analysis (PCoA) showing the impact of stem rot on the rhizosphere fungal community of berseem. Color indicates different sampling sites and symbols indicate healthy and diseased plants. Dissimilarity distances were based on Bray-Curtis distance algorithm. Healthy berseem rhizosphere bacterial microbiome is dissimilar to an infected berseem rhizosphere microbiome.

### Stem rot influences the occurrence and abundance of rhizosphere microbial diversity

The OTUs from all the rhizosphere soil were assigned to 13 bacterial phyla. Microbial diversity at the phylum level showed substantial differences among the rhizosphere of healthy berseem plants and infected berseem plants. The three selected geographical sites demonstrated variable but dependable bacterial phylum share in the rhizosphere microbiome. Proteobacteria was the most abundant bacterial phyla in both infected and healthy Berseem rhizosphere. Its share in total bacterial population in both types of rhizosphere was significantly different such as 88.81- 90.48% for healthy plants and 87.56-89.56% for infected plants. Firmicutes were more abundant in healthy (4.75-6.82%) than infected plant rhizosphere. Members of Bacteroidetes (3.49- 3.65%), Actinobacteria (1.01-1.23%), Cyanobacteria (0.24-0.33%) were dominant in infected plant rhizosphere. Sequences related to Verrucomicrobia, Planctomycetes, Acidobacteria, Armatimonadetes, Gemmatimonadetes, and Chlamydiae were relatively less abundant but detected on both infected and healthy Berseem rhizosphere (**Fig. 3A**).

**Fig. 3.**
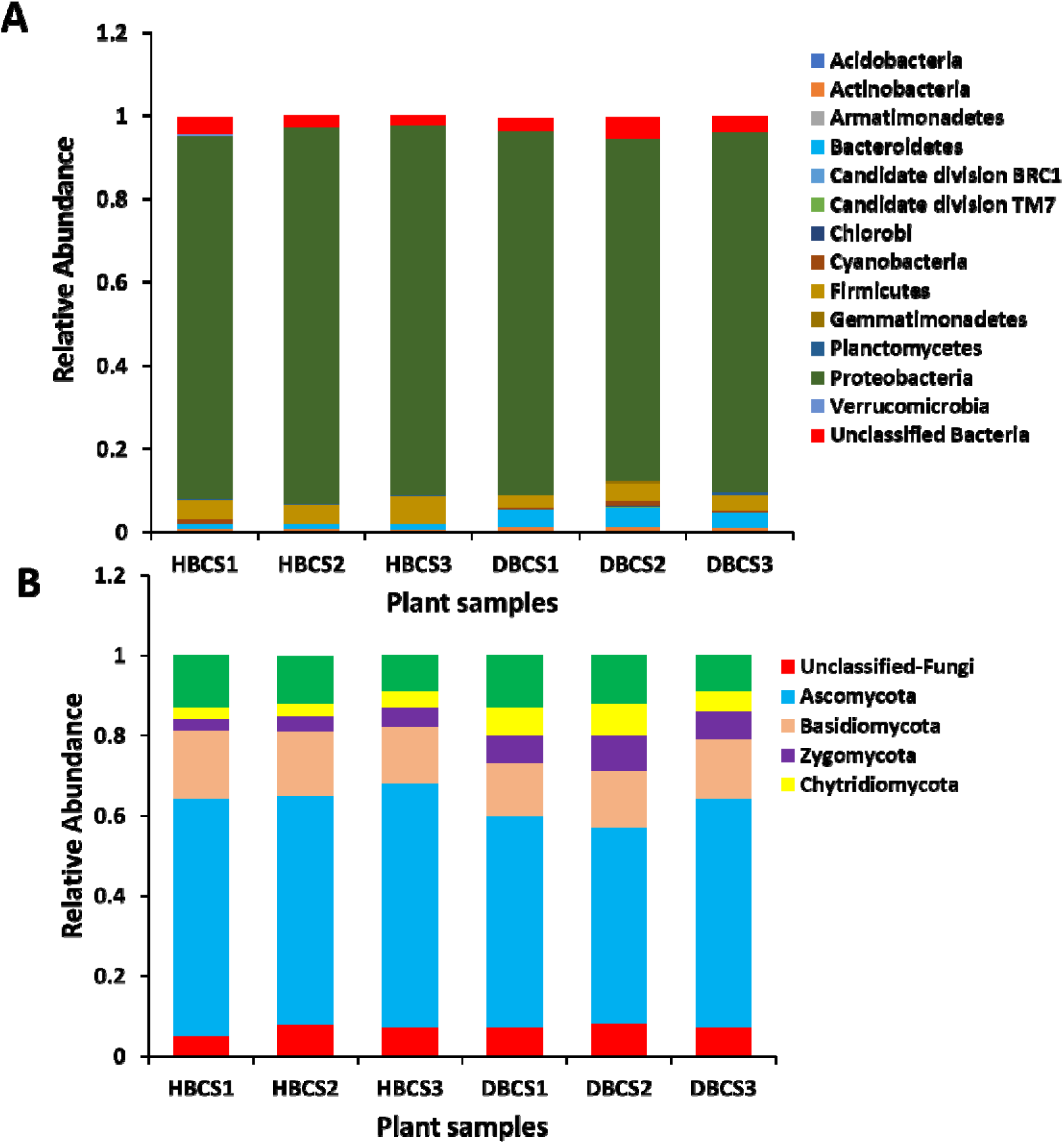
Microbial community composition of healthy and infected berseem at the phylum level. **(A)** Bacterial taxonomic community composition at phylum level, **(B)** Fungal taxonomic community composition at phylum level. Bacterial phyla making up less than 1% of the relative abundance in samples were grouped as Other.

The fungal microbiome was analyzed from infected and healthy Berseem rhizosphere using ITS gene sequences. The ITS sequences were assigned to four fungal phyla predominantly Ascomycota followed by Basidiomycota. As hypothesized significant differences were observed regarding fungal phyla composition in infected rhizosphere and healthy rhizosphere of Berseem plants. Ascomycota (59%) and Basidiomycota (16%) were more abundant fungal phyla in healthy rhizosphere while Zygomycota (9%) and Chytridiomycota (%) were more abundant in infected rhizosphere (**Fig. 3B**).

### Microbial diversity comparison of healthy and infected berseem rhizosphere across geographical sites

The number of OTUs identified from healthy and infected rhizosphere were compared across three selected geographical sites and described by Venn diagram analysis. Overall, maximum bacterial diversity was detected in the infected rhizosphere as 963 bacterial OTUs were detected from three infected rhizosphere sites while 879 bacterial OTUs were detected from healthy rhizosphere soil. A total of 28 common OTUs were identified from all three infected rhizosphere sites, whereas 143, 148, and 154 OTUs were exclusively identified from site 1, 2 and 3, respectively (**Fig. 4A**). Comparatively, 21 common OTUs were identified from all the three healthy rhizosphere sites while 139, 147 and 140 OTUs were distinct within each selected site 1, 2 and 3, respectively (**Fig. 4B**).

**Fig. 4.**
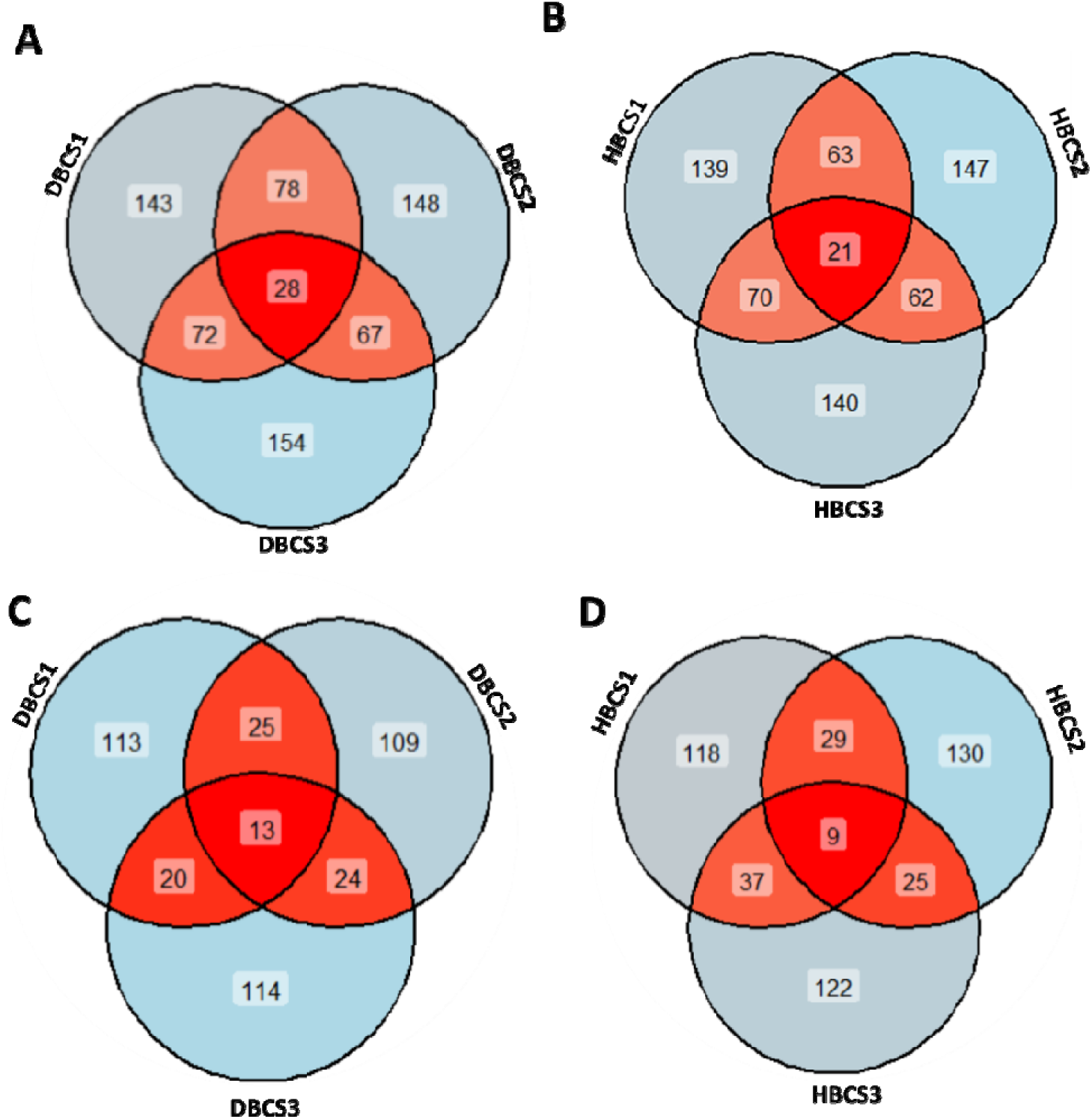
Comparative analysis of bacterial and fungal OTUs from healthy and infected berseem. **A and B** showed the Venn-diagram analysis for bacterial OTUs**. C and D** showed the fungal OTUs comparison

The ITS sequence analysis identified 579 fungal OTUs from healthy rhizosphere of Berseem plant and 513 fungal OTUs from infected rhizosphere. In total identified, 13 fungal OTUs were common in three different infected Berseem rhizosphere geographical sites however 113, 109 and 114 OTUs were exclusive to site 1, 2 and 3, respectively (**Fig. 4C**). In contrast to this, only 9 fungal OTUs were common among the three different geographical sites while 118, 130 and 122 fungal OTUs were exclusive to each selected sampling site (1, 2 and 3, respectively) of healthy rhizosphere of Berseem plants (**Fig. 4D**).

The results for identified bacterial genera between healthy and infected Berseem rhizosphere revealed distinctive pattern irrespective of the geographical locations. A total of 20 characteristic bacterial genera were identified in all three geographical sites with similar richness pattern. The most abundant bacterial genus was *Stenotrophomonas* in both rhizospheres but comparatively more in infected rhizosphere as shown in heat map analysis (**Fig. 5A**). After *Stenotrophomonas*, *Pseudomonas* genus was abundant in both rhizospheres but relatively more in healthy Berseem rhizosphere. *Rhizobium* and *Comamonas* were abundant in healthy plant rhizosphere while *Pantoea* were more abundant in infected plant rhizosphere rhizosphere (**Fig. 5A**).

**Fig. 5.**
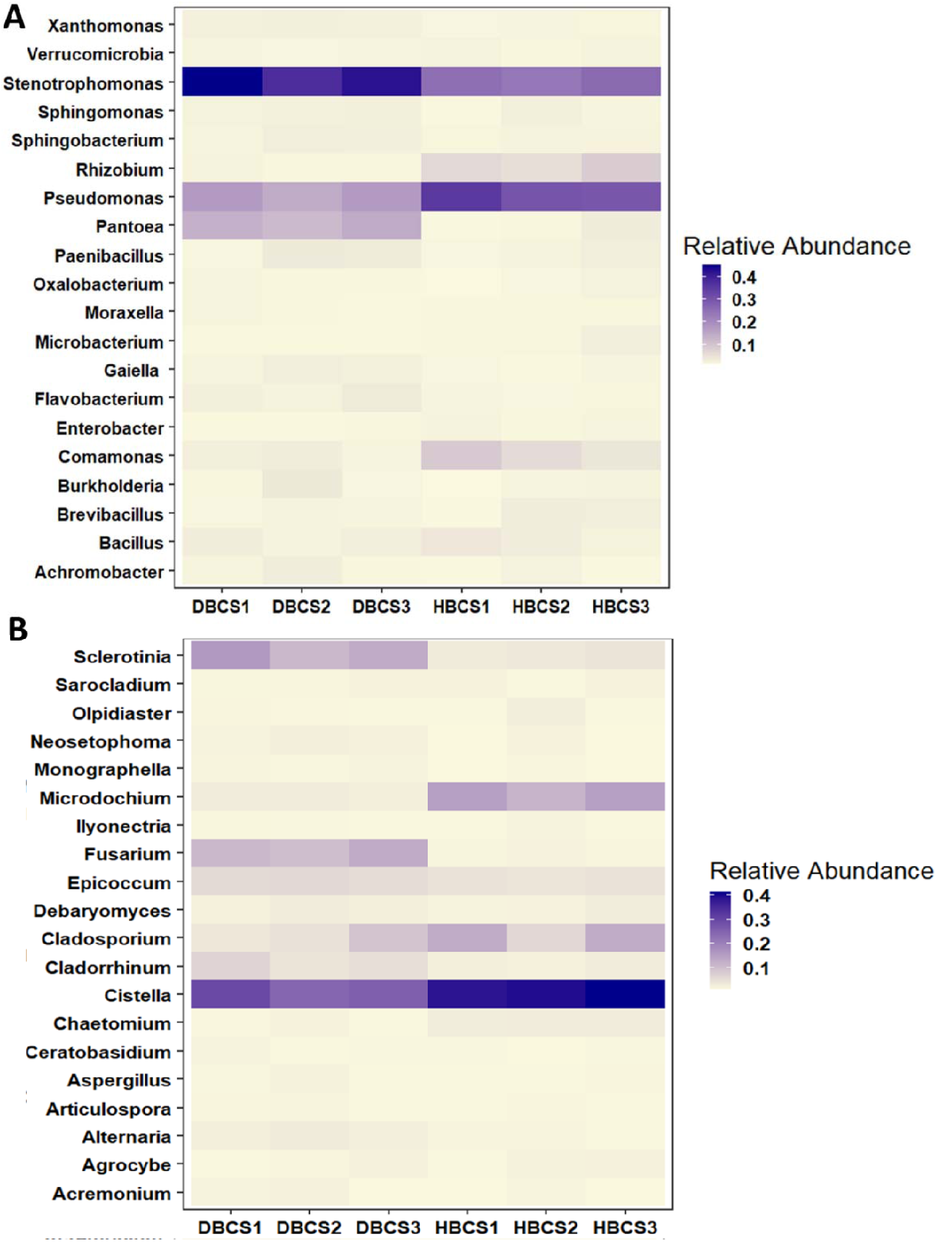
Microbial community composition of healthy and infected berseem at the genus level. **(A)** showed the heatmap for bacterial genera. **(B)** showed the fungal genera comparison

In total, 20 fungal genera were identified from healthy and infected Berseem rhizosphere from each three geographical locations. *Cistella* was the most abundant genus in both rhizospheres but comparatively more abundant in healthy plants than infected plants. *Sclerotinia*, *Fusarium* and *Cladorrhinum* were more abundant in infected plants while *Microdochium* and *Cladosporium* were distinctively abundant in healthy Berseem rhizosphere (**Fig. 5B**).

### Stem rot affects bacterial and fungal functional characteristics

The PICRUSt software based on KEGG pathway database was used to analyze bacterial community function. Bacterial communities from healthy plants’ rhizosphere showed a difference in function prediction as compared to infected plants’ rhizosphere. Most of the functions did not show a significant difference between healthy and infected plants’ rhizosphere. But bacterial communities from healthy plants showed more abundance of bacteria with some selected functions including carbohydrate and energy metabolism, membrane transport, and translation while bacterial communities from infected plants showed more abundance of bacteria with replication and repair, biosynthesis of other secondary metabolites, enzyme families and xenobiotics biodegradation and metabolism (**Fig. 6A**).

**Fig. 6.**
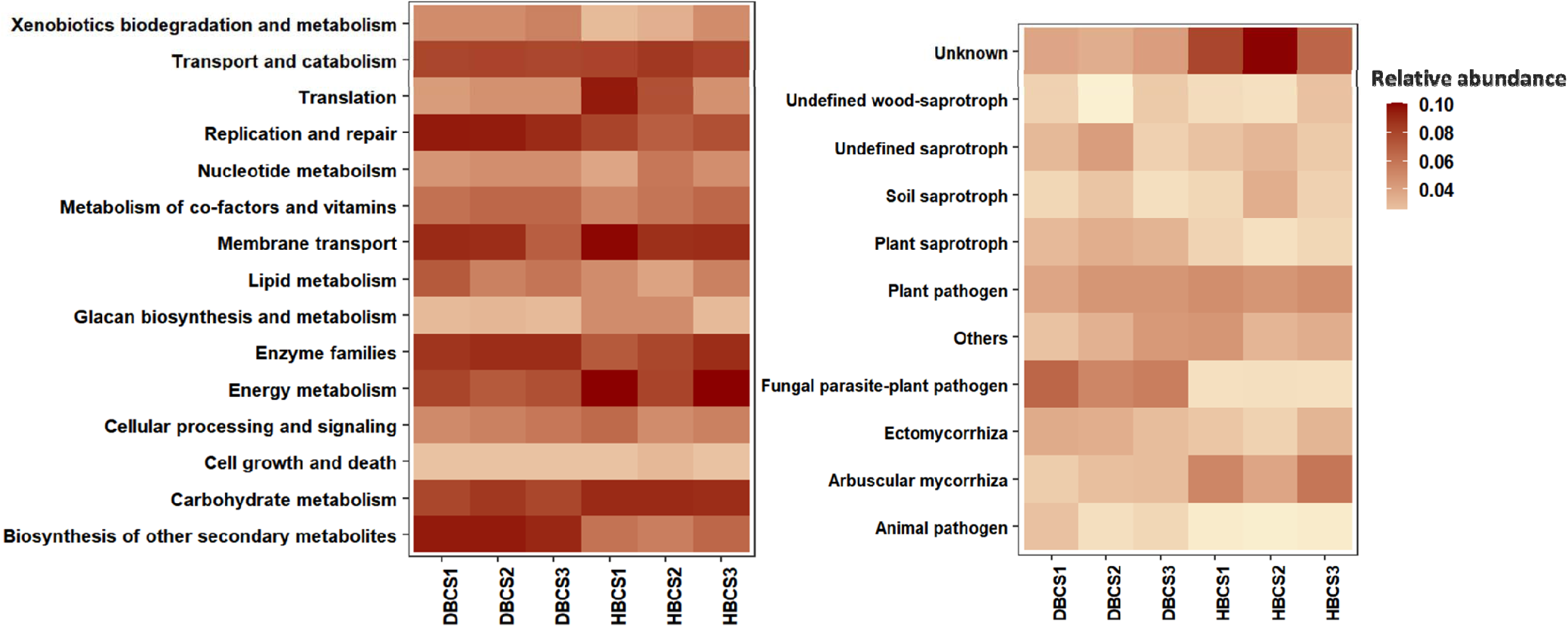
Predicted ecological functions of bacterial and fungal communities between the healthy and infected samples. **(A)** The PICRUSt analysis for function prediction of bacterial communities using KEGG level 2 and **(B)** Functional Guild analysis for function prediction of the fungal communities

Functional Guild (FUNGuild) is a tool used for the functional classification and analysis of fungal communities in healthy and infected plants’ rhizosphere. Functional groups in fungal communities were classified as the fungal parasite-plant pathogen, plant pathogen, undefined saprotroph, plant saprotroph, soil saprotroph, arbuscular mycorrhiza, ectomycorrhiza, undefined wood-saprotroph and animal pathogen (**Fig. 6B**). Some fungal groups such as arbuscular mycorrhiza and soil saprotrophs are decreased in infected plants but plant saprotrophs and fungal parasite-plant pathogens were increased in the infected samples (**Fig. 6B**).

## Discussion

Rhizosphere is a complex environment where microorganisms interact with plant roots and ultimately affect the plant health and productivity (Banerjee et al., 2018; Karuppiah et al., 2020). Microbial diversity in the rhizosphere is considered an important factor that influences plant health and helps to suppress various bacterial and fungal diseases (Miao et al., 2016; Agyekum et al. 2023). At the flowering stage, maximum microbial diversity was observed as compared to the later stage of plant development. Plant roots produce various compounds, especially secondary metabolites to stimulate or inhibit the growth of microorganisms in the rhizosphere (Qiao et al., 2017; Zhalnina et al., 2018; Mukhtar et al. 2020). The relationship between roots and soil microbial diversity plays an important role in plant health and disease resistance. In the current study, we evaluated the response of rhizosphere microorganisms by comparative analysis of healthy and infected plants. We analyzed the bacterial diversity using 16S rRNA gene and fungal diversity using ITS gene from the rhizosphere of healthy and stem rot infect berseem clover. To the best of our knowledge, this study is the first report on the response of the rhizosphere microbiome of berseem clover to stem rot infection.

Microbiome analysis showed a significant decrease in both bacterial and fungal diversity in the infected berseem rhizosphere as compared to the healthy berseem rhizosphere. Alpha diversity indices including observed OTUs, Shannon index and FaithPD were higher in healthy plants as compared to infect ones across all the sampling sites (**Fig 1**). Beta diversity analysis also revealed that microbial communities from the infected plant rhizosphere clustered separately from microbial populations from healthy plant rhizosphere (**Fig 2**). Some previous studies such as Wang et al. (2021) reported a significant decrease in overall microbial populations in the rhizosphere of rotten Sanqi ginseng and an increase in the abundance of potentially pathogenic fungi. Another study also reported an abundance of certain bacterial and fungal taxa such as *Microccocus, Ralstonia, Fusarium, Ilyonectria,* and *Plectosphaerella* in the rhizosphere of avocados afflicted by the *Phytophthora* root rot (Solís-García et al. 2020). Root rot induces the proliferation of specific microbial populations that produce antimicrobial compounds to suppress or resist disease (Pascale et al. 2020).

The microbiome analysis showed that the rhizosphere’s bacterial and fungal community changed significantly between healthy and diseased soils (**Fig. 3A and 3B**). We further found noticeable differences among the richness of Proteobacteria and Firmicutes in the rhizosphere of healthy and diseased plants (Hayden et al., 2018). The reduction in this richness indicates the dominant interspecies behavior of stem rot fungus in diseased plant rhizosphere. This microbial richness is consistent in heatmap analysis where the genus *Stenotrophomonas* is found in higher abundance than other identified bacterial genera in the diseased rhizosphere (**Fig. 5A**). This shift in abundance is an indication of increased plant defense due to the plant growth promotion properties of *Stenotrophomonas* as reported in a similar context where the diseased suppressive rhizosphere has a higher abundance of this genera (Ling et al., 2024; Sharma et al., 2024). *Stenotrophomonas* are naturally found in the rhizosphere of many plants and are documented for their antifungal properties and providing biocontrol to the host plant. Similarly, this was confirmed in our findings where the healthy rhizosphere has a lower abundance as compared to the diseased rhizosphere of the Berseem plant (Ciftci et al., 2023).

Comparing the relative abundance of fungal phyla, we observed that the abundance of Ascomycota reduced while the abundance of Basidiomycota, Zygomycota and Chytridiomycota increased in the diseased rhizosphere. We infer that the reduction of Ascomycota richness in the diseased rhizosphere may be due to the dominant behavior of stem rot fungus in interspecies competition (Li et al., 2022) and synergistic behavior among pathogenic fungi and members of Basidiomycota, Zygomycota and Chytridiomycota phylum. Similar to our findings another study by Dubey et al. (2023) reported a clear shift of fungal communities and *Ascomycota* remained the most abundant phyla followed by *Mortierellomycota*, *Chytridiomycota*. The dominant fungal genus was *Cistella* in both rhizosphere but relatively its abundance is reduced in diseased rhizosphere in comparison to the healthy soil. A more prominent increase in the fungal genus was observed for *Sclerotinia*, *Fusarium*, *Epicoccum*, and *Cladorrhinum* in the diseased rhizosphere. While increased abundance of *Microdochium Cladosporium* and *Chaetomium* fungal genus was observed for a healthy rhizosphere.

The functional gene analysis demonstrated that diseased rhizosphere had a major shift for increased bacterial secondary metabolites production followed by replication and repair, enzyme family and xenobiotics biodegradation and metabolism gene expressions. While reduced gene expression was observed for translation, glycan biosynthesis and metabolism, cellular processing and signaling (**Fig. 6A**). Finally, we analyzed the fungal gene functions associated with the disease and healthy rhizosphere using the OUT data. The functional annotation categorized fungal communities into ten functional categories and the heatmap (**Fig. 6B**) showed variable gene expression. We observed increased functional gene expression in diseased rhizosphere for unidentified saprotrophs, plant saprotrophs, fungal parasite-plant pathogens, and animal pathogens. Similar to our findings another study (Duan et al., 2023) reported seven categories of fungal functional annotations dominated by saprotroph, pathotroph-saprotroph-symbiotroph, pathotroph, symbiotroph. Our study provides new insights into the dynamics of the rhizosphere microbiomes of diseased and healthy berseem plants for future biocontrol and fertilization regimes.

## Conclusion

The present study showed the effect of stem rot on the bacterial and fungal communities associated with berseem clover rhizosphere. Microbiome analysis revealed that bacterial and fungal communities associated with healthy plants are significantly different from microbial communities associated with stem rot-infected plants. Members of Proteobacteria and Firmicutes were more dominant in healthy plants while Bacteroidetes and Actinobacteria were more dominant in infected plants. Bacterial genera *Rhizobium* and *Comamonas* were more abundant in healthy plants while *Pantoea* was more abundant in infected plants and fungal genera *Sclerotinia*, *Fusarium* and *Cladorrhinum* were more abundant in infected plants while *Microdochium* and *Cladosporium* were distinctively abundant in healthy Berseem. Functional characterization of microbial communities described that bacterial communities from infected plants had more abundance of bacteria with functions replication and repair, enzyme families, and biosynthesis of other secondary metabolites as compared to healthy plant microbiome and decreased in fungal groups including arbuscular mycorrhiza and soil saprotrophs and an increase in plant saprotrophs and fungal parasite-plant pathogens. This study could help to understand how the microbiome reacts to biotic stress such as fungal diseases and to understand how the microbiome could help plants induce resistance against fungal diseases.

---

**Figure S1.**
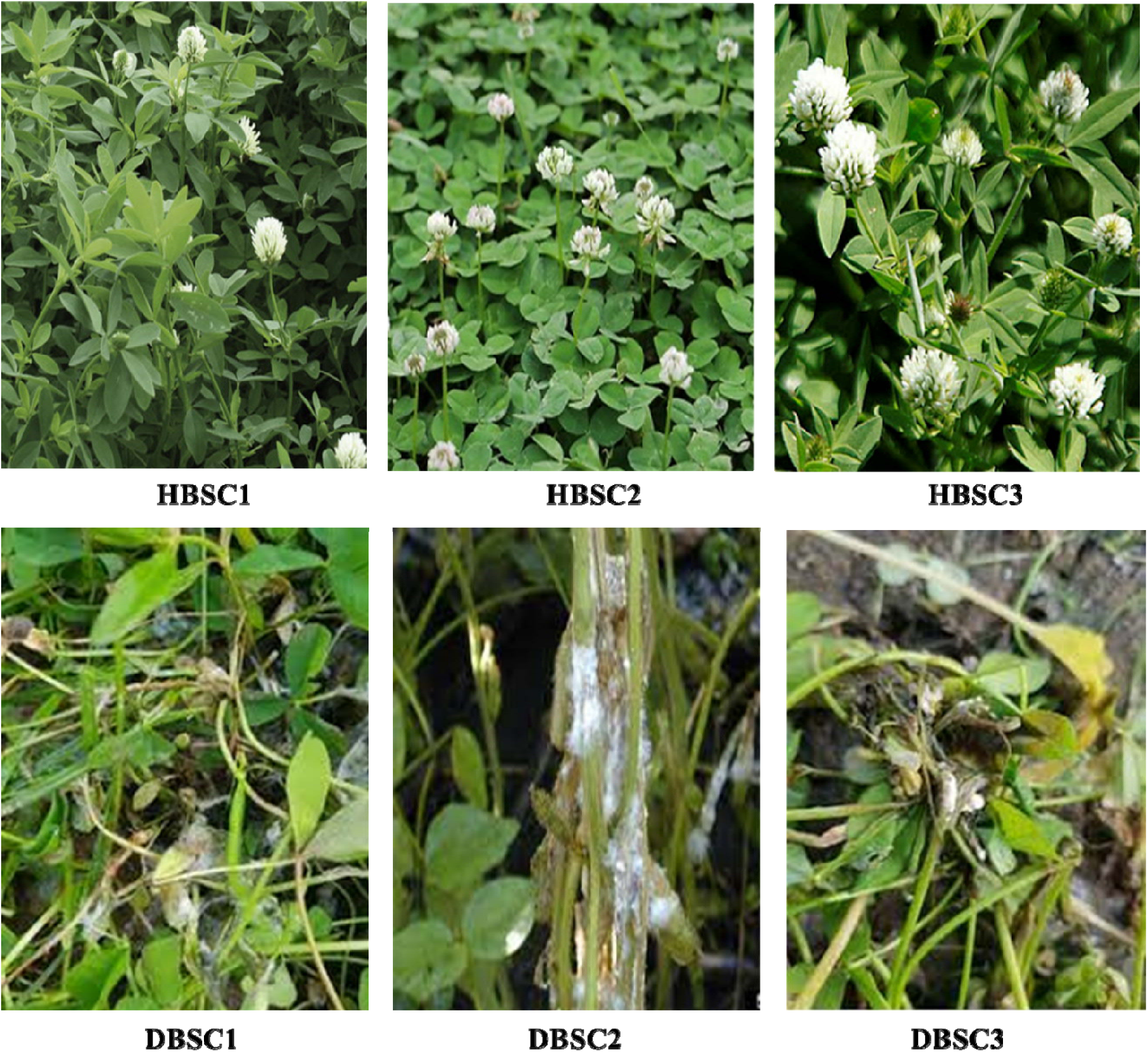
Sampling of healthy and stem rot infected plants from different locations.

